# Spatial constraints subvert microbial arms race

**DOI:** 10.1101/2023.06.16.545151

**Authors:** Raymond Copeland, Peter J. Yunker

## Abstract

Biofilms, surface attached communities of microbes, grow in a wide variety of environments. Often, the size of these microbial community is constrained by their physical surroundings. However, little is known about how size constraints of a colony impact the outcome of microbial competitions. Here, we use individual-based models to simulate contact killing between two bacterial strains with different killing rates in a wide range of community sizes. We found that community size has a substantial impact on outcomes; in fact, in some competitions the identity of the most fit strain differs in large and small environments. Specifically, when at a numerical disadvantage, the strain with the slow killing rate is more successful in smaller environments than in large environments. The improved performance in small spaces comes from finite size effects; stochastic fluctuations in the initial relative abundance of each strain in small environments lead to dramatically different outcomes. However, when the slow killing strain has a numerical advantage, it performs better in large spaces than in small spaces, where stochastic fluctuations now aid the fast killing strain in small communities.

Finally, we experimentally validate these results by confining contact killing strains of *Vibrio cholerae* in transmission electron microscopy grids. The outcomes of these experiments are consistent with our simulations. When rare, the slow killing strain does better in small environments; when common, the slow killing strain does better in large environments. Together, this work demonstrates that finite size effects can substantially modify antagonistic competitions, suggesting that colony size may, at least in part, subvert the microbial arms race.

**Author summary:** Biofilms are often crowded with many bacteria in direct contact. As a result, the competition for space and resources often turns deadly. Bacteria have evolved many mechanisms with which to kill each other; this bacterial warfare is often studied in large communities on agar plates or in flow cells [1]. However, in nature these colonies are often smaller, due to spatial constraints or shear forces. It is unclear how bacterial warfare proceeds in small systems.

We performed individual based model simulations of bacterial warfare comprising two strains, each capable of killing the other on direct contact. We found that the community size played a substantial role in determining the outcome. When at a numerical disadvantage, the slow killing strain survived at much higher rates in small communities. In fact, there were many conditions in which the slow killing strain survives in small spaces but is completely eliminated in large ones. Conversely, when the slow killing strain is more common, it performs better in large spaces. Together, these observations demonstrate that finite size effects aid the strain that is at a disadvantage, and in some conditions, can even flip which strain increases its abundance.

Finally, we experimentally tested the results of these simulations. Two mutual killing strains of *V. cholerae* were grown unconfined on agar plates (i.e., in large spaces) or confined within square holes with sides 7.5*μm* long (i.e., in small spaces). In these experiments we found that the slow killing strain survived at significantly higher rates in confinement, validating simulation results.

## 1 Introduction

Bacteria in nature are often in close contact and form biofilms, crowded, surface attached communities by secretions of an extra cellular matrix [2]. These structures comprise a multitude of different species and strains [3], and thus social interactions are commonplace. These can be direct, such as quorum sensing [4], or indirect, but broadly, these interactions can be categorized based the strains aiding or inhibiting one another: mutualism, parasitism, competition, etc [5]. Competition, where strains inhibit one another strain, often includes antagonistic interactions [1, 6]. As a result, fitness is often a function of killing ability, potentially initiating a microbial arms race [7]. For example, the Type VI secretion system (T6SS), a contact dependent killing mechanism, is found in 25% of gram negative bacteria [8]. However, the effectiveness of the T6SS has been found to depend sensitively on context [9–11], and in many cases the T6SS is not necessary for bacteria to colonize occupied terrain [12, 13]. While it is clear that microbes do not experience a run-away arms race, it is less clear how this race has been subverted.

Spatial structure is known to play an important role in antagonistic competitions [9, 14–18]. Further, environmental structures, such as surface roughness or anisotropic height perturbations, within large communities are known to impact competitive outcomes, and even stabilize coexistence [19, 20].Crucially, in nature colonies are often spatially constrained by their environment, limiting how large they can grow. These size constraints arise from many different sources, such as environmental barriers and external stresses [21–24]. However, experimental studies of microbial antagonism are typically performed on agar plates or in flow cells, which allow colonies to grow to large sizes, containing thousands of cells or more.It has been previously shown that spatially limited and fragmented cells exhibit different social dynamics and prey cells survive at higher rates in these environments compared to large ones [17, 25] However, little is known about how antagonistic competitions proceed in small environments and with small numbers of cells, despite the natural relevance of such settings.

Here, we use agent-based simulations to explore the effects of spatial constraints on bacterial contact killing, and then validate our observations experimentally. In simulations of mutual killing strains, we find that community size can substantially impact the outcome. When it lacks a numerical advantage over a faster killing strain, the slower killing strain survives at much higher rates in smaller communities than in larger communities. We show that this effect arises from fluctuations in initial compositions, which are more substantial in small communities. Further, when the slow killing strain has a numerical advantage over a fast killing strain, the slow killing strain is more successful in large communities than in small communities, where finite size effects now benefit the fast killing strain. The outcome of contact killing competitions thus depends on community size. We test these ideas experimentally by inoculating two mutual-killing strains on transmission electron microscopy (TEM) grids, confining bacteria to ∼ 50*μm*^2^ holes. Antagonistic competitions in TEM grids are highly spatially constrained; we compare these experiments to colonies grown unconfined on agar plates (∼ 10*E*6*μm*^2^). As in our simulations, we experimentally found that the outcome depends both on the community size and initial abundances. When the slow killer is at an initial numerical disadvantage, it performs better in the confined environment; when the slow killer is at an initial numerical advantage it performs better on bare agar.

Thus, experiments also demonstrate that community size impacts contact killing competition outcomes.

## 2 Material and Methods

### 2.1 Simulation Methods

We investigated the impact of community size on contact killing competitions with agent based simulations. We simulate individual cells pushing, growing, reproducing, and killing over time. Our simulations all feature two strains that kill each other on contact; we vary the kill rate of the slow killer to isolate the benefit of contact killing superiority. We perform simulations for many different community sizes, to determine the impact of size on warfare outcomes.

We modeled cells as interacting via a simple harmonic repulsive potential:

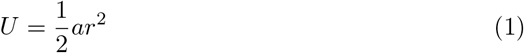

where *a* = 500 is the harmonic potential constant and *r* is the radial overlap between two neighboring cells (Fig. 1 b). The motion of cells is over-damped, i.e., cells only move when actively experiencing a net force. This approach was inspired by previous work that found that bacterial interactions in biofilms are accurately modeled with stiff, longer ranged interaction potentials with over-damped dynamics [26, 27]. At the beginning of each time step, the order in which cells interact is set at random; as we iterate through the list, we update each cell’s position as overlapping neighbors push the focal cell.

**Fig 1.**
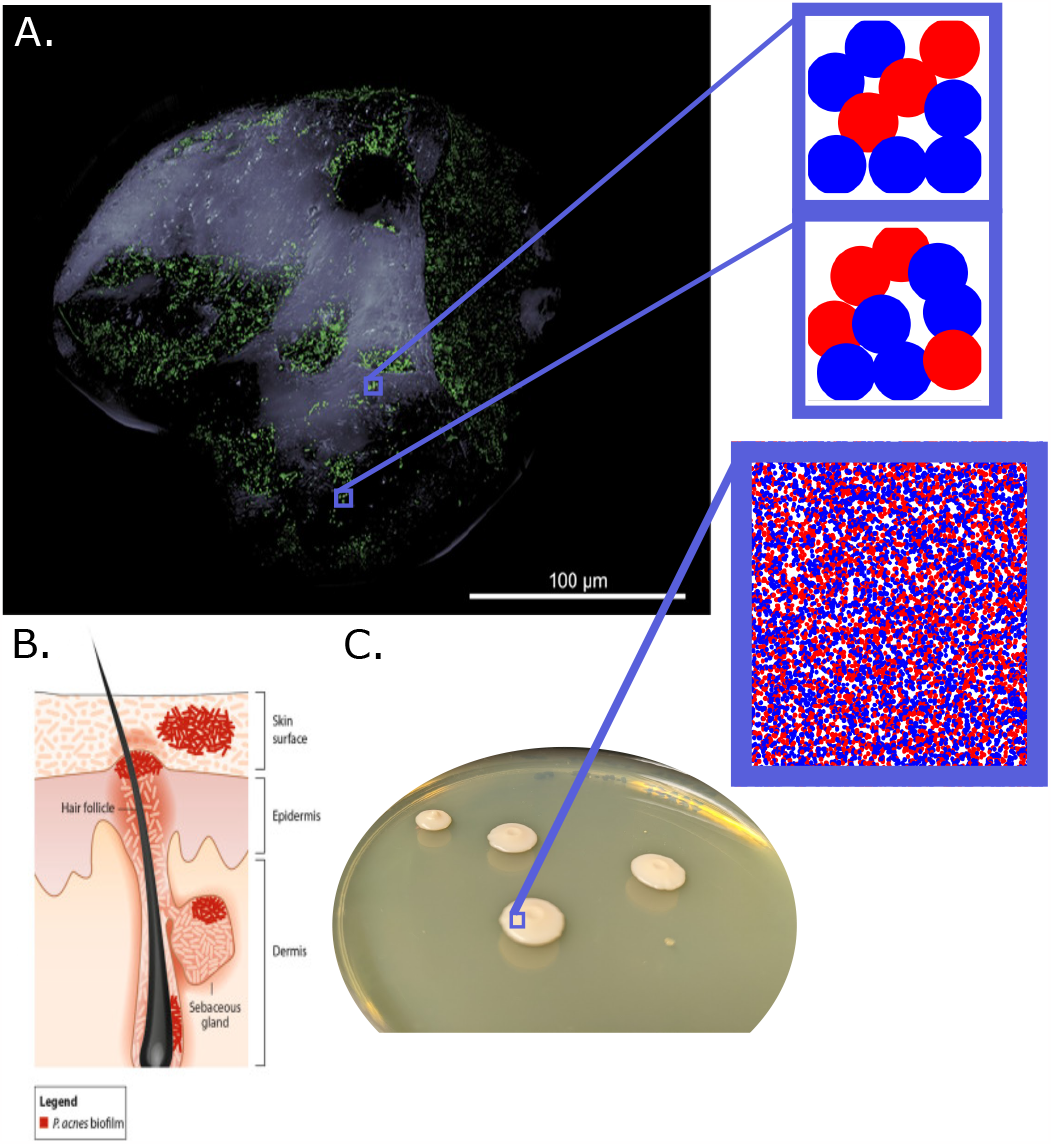
Microbial communities in nature are often size constrained. A. Microbial colonies studied in the lab are often large, such as these *V. cholerae* colonies grown on agar. However, colonies in nature are often much smaller. For example, biofilms in skin pores (B, adapted from [21]) and on grains of sand (A, adapted from [23]) are significantly smaller than those studied in the lab. A and B were used under the Creative Commons Attribution 4.0 International License. C. In our simulations we confine two mutually antagonistic strains (shown in red and blue) to systems of different sizes but with similar cell densities. We show simulations of *N*_*max*_ = 9 cell systems and a *N*_*max*_ = 1000 cell simulation with approximate scaling for context. The circles show the extent of the soft intercellular interaction potential.

Each cell begins with a radius, *R*, of 0.5. Cell mass grows at a constant rate to simulate a homogeneous nutrient distribution, and to isolate the effect of contact killing, rather than growth dynamics and competition for nutrients, on the outcome [1]. We thus model the change in radius per unit time as:

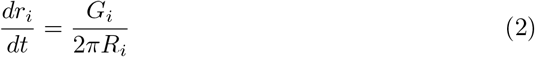

where *G*_*i*_ is the mass growth rate. This approach produces an area that increases linearly with time (i.e., mass increases at a constant rate). Cell division occurs when a cell has doubled in mass, the mother cell spontaneously splits in place into two daughter cells each with half of the maximum mass and with the minimum radius *R*_*min*_. To prevent synchronized cellular division across the community, we assign values of *G*_*i*_ to each cell randomly from a uniform distribution with coefficient of variation 0.1 around the mean, 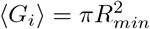 [27, 28]. This process takes on average Δ*t* = 1.0 (one generation) from when an individual cell starts growing to when it divides. To emulate experimental observations that microbial growth stops under sufficiently large mechanical stresses [29], cell growth in our simulations stops if the system-wide packing fraction exceeds a threshold, *ϕ*_*t*_, determined by the system size. *ϕ*_*t*_ is determined by:

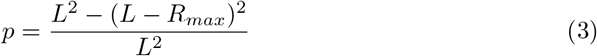

and

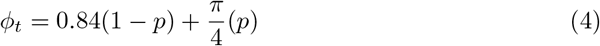

where L is the length of the system size. These equations weigh the known packing fraction (0.84) that characterizes bulk jammed disks in two-dimensions [30, 31] with the highest packing fraction a disk can reach on a hard edge (*π/*4). *p* is the proportion of space beyond which the largest cells can interact with the edge, which varies with system size.

Focal cells can potentially kill a cell of the opposing strain if their interaction radii overlap, i.e., if they are in contact. Each strain is assigned a distinct killing rate in units of killing events per time per cell. We will refer to the strain with the lower killing rate as the slow killing strain and the strain with the higher killing rate as the fast killing strain. In all simulations, we hold the killing rate of the fast killing strain, *κ*_*F*_, at *κ*_*F*_ = 1, i.e., on average each of the fast killing strain takes 1 generation to kill an opposing cell (a rate in line with previous works [32]). We vary the killing rate of the slow killing strain (*κ*_*S*_, which is always ≤ 1), enabling us to characterize these competitions by their ratio (*κ*_*r*_ = *κ*_*F*_ */κ*_*S*_).

The killing probability per time step is the rate multiplied by the length of the time step (*dt* = 0.01), which produces an exponentially distributed “time to kill.” If the focal cell contacts its opposing strain, then it may attempt to kill. If the focal cell overlaps with multiple opposing strain cells, one of the opponents is randomly selected for the focal cell to attempt to kill, and the focal cell is allowed to attempt to kill once, regardless of the number of opposing cells that overlap with it. After selecting a target, we generate a random number between 0 and 1; if the random number is smaller than the strain’s killing probability, then the focal cell kills the target cell and the target cell disappears.

Cells interact within square environments with varying size. These environments are bounded by stiff walls, modeled with a harmonic repulsive potential with a harmonic potential constant twice that of the cells themselves, thus ensuring cells remain well-confined. We vary the length, *L*, of simulated regions (*L* = 3, 5, 8, 10, 31, and 100), and we then find the maximum packing fraction of each region according to equations 3 and 4 (*ϕ* = 0.817, 0.826, 0.831, 0.833, 0.838, 0.839 for *L* = 3, 5, 8, 10, 31, and 100, respectively) and then fill the region with the maximum number of cells, *N*_*max*_, of minimum radius (0.5) to start a simulation (*N*_*max*_ = 9, 26, 68, 106, 1025, 10686 cells for *L* = 3, 5, 8, 10, 31, and 100, respectively). We place cells into their environment with random positions. To simulate cells randomly attaching to a surface from a planktonic suspension, the proportion of each strain in a given community is determined stochastically. We set the fraction of the slow killing strain in the planktonic suspension, *p*_*i,S*_. Upon inserting each cell into the environment, we call a random number between 0 and 1; if it is less than *p*_*i,S*_ the cell is set as the slow killing strain, otherwise the cell is set as the fast killing strain. From this stochastic seeding, the initial abundance of slow killers *S*_*i*_ varies from simulation to simulation. We run each simulation for 6400 time steps, i.e., 64 generations, ensuring that all simulations reach a steady state wherein the relative abundance of the two strains changes by less than 0.1%, i.e., no longer significantly changes, over one generation. For each community size (*L*), we simulate many replicates and aggregate the competition results to form a meta-community of many *L* sized environments.

### 2.2 Experimental Methods

We used two strains of *Vibrio cholerae* [33] that use the Type VI Secretion System (T6SS) [34] to deliver toxins to neighboring cells in direct contact. The two strains are isogenic except for their toxins, anti-toxins, and fluorescent proteins. Both strains express Green Fluorescent Protein GFP, but only the slow killing strain expresses Red Fluorescent Protein RFP. In all experiments, the two strains were grown separately in LB broth for 24 hours and then mixed together. They were then inoculated with 2*μl* drops on open agar. The slow killing strain was identified by mixing both strains at the same optical density and inoculating on an agar pad; after 24 hours of growth we identified the slow killing strain as the strain that occupied a smaller proportion of space. To confine this competition to small system sizes, we also inoculated bacteria on top of transmission electron microscopy (TEM) grids that were placed on top of LB agar. All steps of all experiments were done at 37^*°*^ C. The grids are 3.05 mm in diameter and have over 2000 square holes, each with length and width of 7.5*μm* (Fig. 7a). Each hole in the grid which can hold ≈ 35 cells in a single layer assuming a high density and using the typical size of a *Vibrio cholerae* [35].

The colonies were imaged on a confocal microscope (Nikon A1R); these measurements returned data for the red, green, and transmitted light channels. Since the two strains differ in fluorescent proteins and toxins, we used confocal microscopy to determine the final proportion of each strain. The proportion of the slow killer was measured in each sample by measuring the number of pixels where the red channel value exceeding the green channel and dividing that number by the total number of pixels that cells occupied; Cells occupied the entire imaged region for the bare agar experiments and only the space in-between the grid lines for the TEM experiments. The transmitted light channel was used to determine which pixels were grid space by binarizing the channel using an Otsu [36] threshold. The pixels that corresponded to the grid were excluded from all calculations.

## 3 Results

### 3.1 Effect of system size on stochastic competitions

To determine the impact of community size on competitions between mutually killing strains of bacteria, we simulated communities in different sized environments (*L*) and strains with different killing ratio *κ*_*r*_ and equal initial abundances (*p*_*i,S*_ = 0.5). We characterize the outcome by the final relative abundance of the slow killing strain 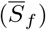, averaged across many simulations, as a function of *κ*_*r*_ (Fig 2). We found that in large spaces (i.e., *N* = 68, 106, 1025, and 10686), there is a critical value for the killing ratio, *κ*_*r*_ ≈ 0.4, below which the slow killer is completely eliminated, i.e., 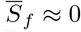. For these large systems, as *κ*_*r*_ increases above 0.5, the slow killing population 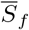 quickly increases to 0.5 when *κ*_*r*_ = 1.0, as expected for strains with equal killing rates and equal initial relative abundances (Fig 2 A). For small environments (*N*_*max*_ = 9 and 26), a different trend emerges; the relative slow killer population 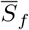 is a linear function of the killing ratio *κ*_*r*_ (*R*^2^ = 0.991 and 0.946, for *N*_*max*_ = 9 and 26, respectively) and 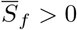 for most values of *κ*_*r*_ we simulated.

**Fig 2.**
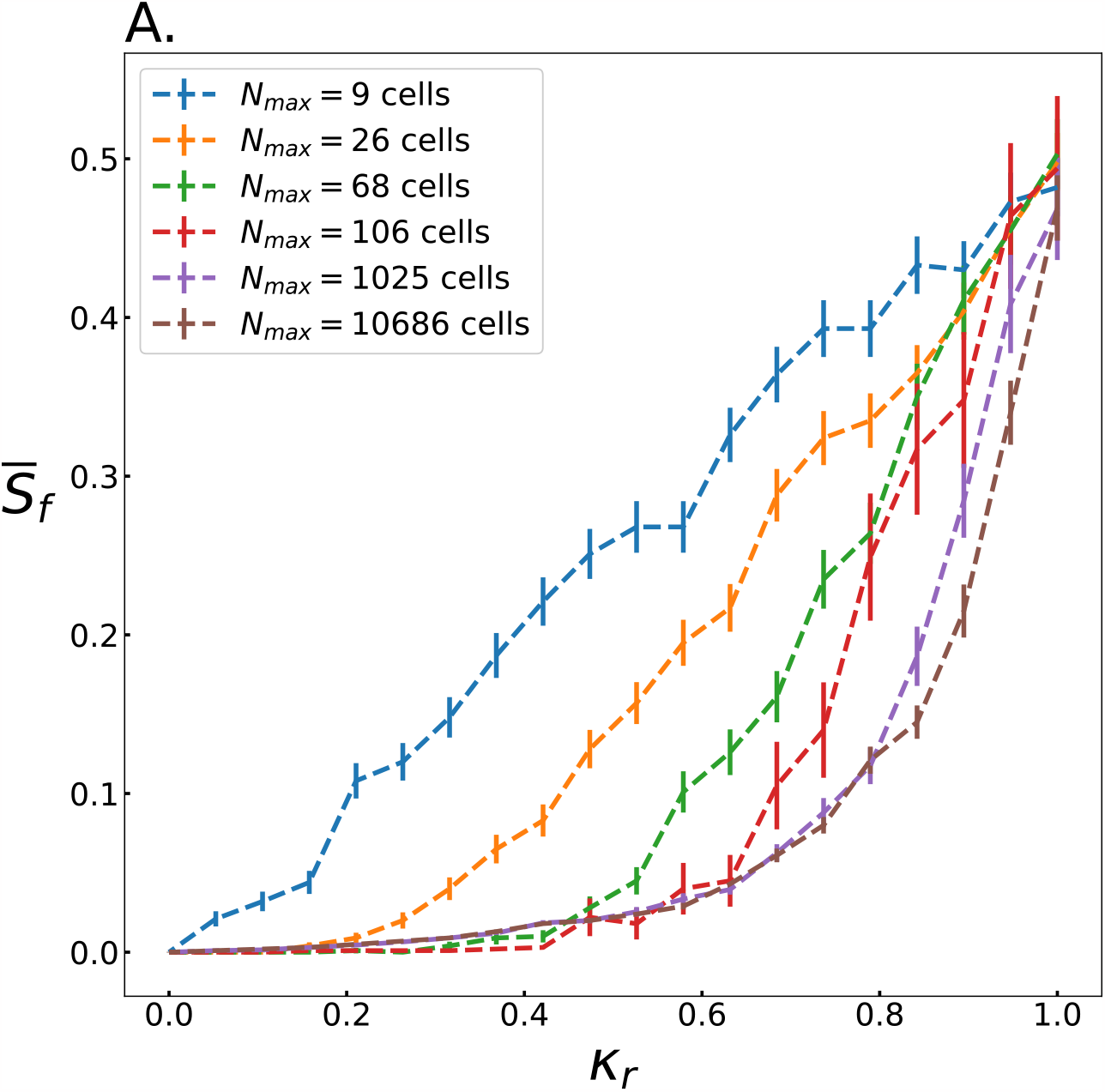
Slow killing strains survive better in small spaces with equal starting abundance. A. The fractional mean relative abundance of slow killing cells, 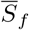, after 64 generations of simulation is plotted against the relative killing rate of the slow killer, *κ*_*r*_, for different size simulations. Each line consists of 20 evenly spaced data points. Each data point is the mean of many simulations; for the *N*_*max*_ = 9, 26, 68, 106, 1025, and 10,686 cell simulations we average across 750, 750, 500, 100, 50 and 20 simulations, respectively. For 106, 1025, and 10,686 cell simulations 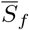 decreases rapidly with decreasing *κ*_*r*_. Conversely, for *N*_*max*_ = 9 cell and 26 cell simulations 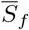 decreases linearly with *κ*_*r*_. The standard error in 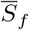 from the simulations are shown along each trend.

To understand the source of the finite size effects observed in figure 2, we next investigate the role of stochastically seeding the initial relative abundance of slow killers (*S*_*i*_) in each simulation. To directly test the effect of *S*_*i*_ on outcome 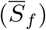, we ran simulations in which *S*_*i*_ is set deterministically rather than stochastically. We simulated a representative selection of killing ratios *κ*_*r*_ and initial abundances *S*_*i*_ for environments with *N*_*max*_ = 9 cells and environments with *N*_*max*_ = 1025 cells, and we measured 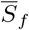 (fig 3 a and fig 3 c). We found that increasing either *S*_*i*_ or *κ*_*r*_ produces an increase in 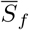, as expected (fig 3 a and c). However, there are more initial conditions that enable the survival of the slow killer in the small system than there are in the large system. Further, individual simulations of the small system typically end with only a single strain surviving, resulting in a large standard deviation in 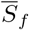 (fig 3 b and fig 3 d). In contrast, individual simulations of large systems are more deterministic in their outcomes, thus producing smaller standard deviations in 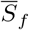 (fig 3 d).

**Fig 3.**
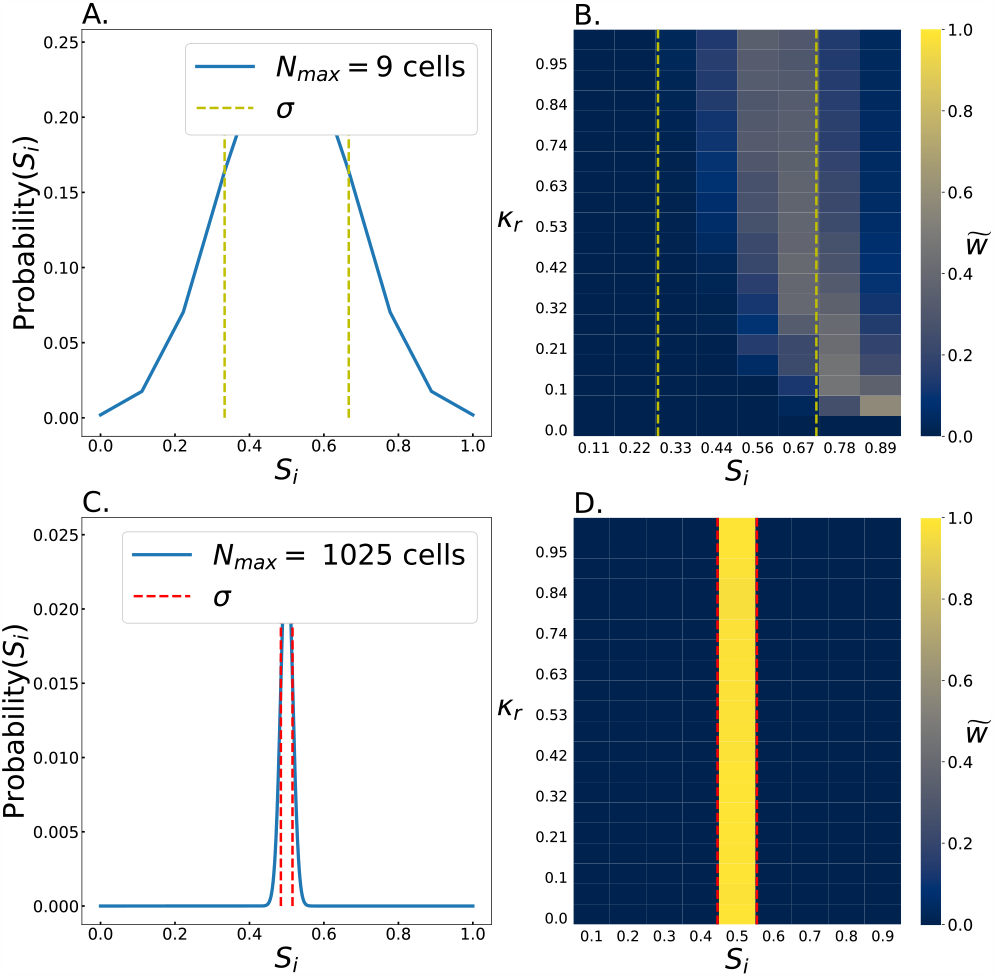
Slow killing cells in small systems survive in more starting conditions. A. This heat map shows how the final relative abundance of the slow killing cells, 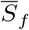, depends on its initial relative abundance, *S*_*i*_, and its relative killing rate, *κ*_*r*_, in *N*_*max*_ = 9 cell competitions. Each data point is the result of 1000 simulations. The yellow trend line shows where 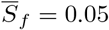. This heat map shows the standard deviation of 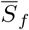 from simulations in A. The standard deviations are often quite large as competition in *N*_*max*_ = 9 cell systems typically ends with one strain being eliminated. C. This heat map shows how the final relative abundance of the slow killing cells, 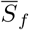, depends on its initial relative abundance, *S*_*i*_, and its relative killing rate, *κ*_*r*_, in *N*_*max*_ = 1025 cell competitions. In contrast to the *N*_*max*_ = 9 cell system outcomes, for *N*_*max*_ = 1025 cell systems, 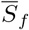 is non-zero for a smaller range of *S*_*i*_ and *κ*_*r*_. Each of the *N*_*max*_ = 1025 cell data points is the result of 49 simulations. The red trend line shows where 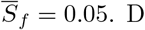. This heat map shows the standard deviation of 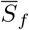 from simulations in C. The standard deviations here are small compared to the *N*_*max*_ = 9 cell simulations.

To understand the source of these fluctuations, we next delineated the distribution of initial conditions in stochastically seeded simulations. We calculated the probability of different *S*_*i*_ values occurring for systems with *N*_*max*_ = 9 cells and systems with *N*_*max*_ = 1025 cells following a binomial distribution. Every simulation of a given system size starts with the same number of cells; the probability of any cell being a slow killer is 0.5. Thus, the probability of different initial numbers of slow killers is:

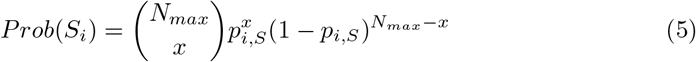

where *N*_*max*_= 9 or 1025, *x* is the initial number of slow killer cells, *S*_*i*_ = *x/N*_*max*_, and *p*_*i,S*_ is 0.5 as set by well mixed, large planktonic suspension (Figs. 4a and c). In small spaces, fluctuations can potentially produce a large initial proportion of slow killers (fig 4 a); in fact, *S*_*i*_ *>* 0.55 50% of the time. Conversely, in large spaces stochastic seeding is unlikely to substantially modify the initial proportion of slow killers compared to the platonic suspension (fig 4 c); in fact, *S*_*i*_ *>* 0.55 only 0.7% of the time.

**Fig 4.**
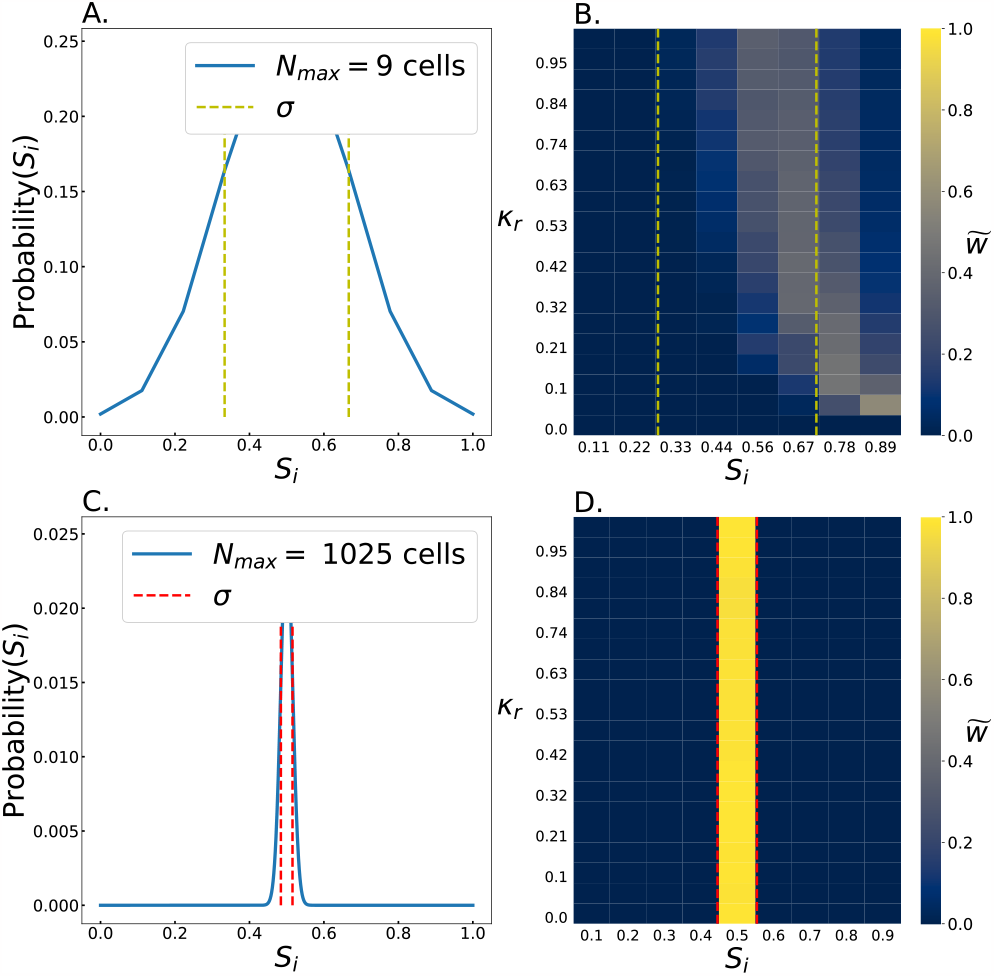
Fluctuations in small systems favor the slow killing strain. The probability distribution of the initial relative abundance of slow killing cells, *S*_*i*_, is shown for *N*_*max*_ = 9 cell and *N*_*max*_ = 1025 cell systems (A and C, respectively) with on standard deviation outlined for each (yellow for small systems and red for large). The distribution of *S*_*i*_ is of course much broader for the *N*_*max*_ = 9 cell system than for the 1000 cell system. B, D. These heat maps show the relative impact of each *S*_*i*_ for determining 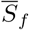, as a function of *κ*_*r*_. This calculation is done by weighing the values of 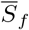 for different *S*_*i*_ from Fig. 3 A and C by the probability that a particular value of *S*_*i*_ occurs (from panels A and C in this figure). This number (*w*) is then normalized across each row, for ease of viewing. In effect, this quantity 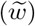 represents how much each *S*_*i*_ contributed to the results seen in Fig. 2 A. The first standard deviation for panels A and C are shown in B and D for each respective environment.

We next quantified how much each stochastically set value of *S*_*i*_ contributed to survival of slow killers (i.e., 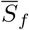). In other words, we accounted for the fact that larger values of *S*_*i*_ are likely to lead to larger values of 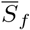, but are progressively less likely to occur. We thus weighed each outcome of the deterministic simulations (where we set *S*_*i*_) by its probability of occurring from equation 5 (fig 4 a and c) to enable comparisons:

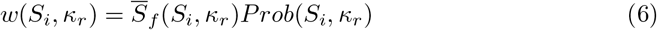

We then normalized these values across rows of constant *κ*_*r*_:

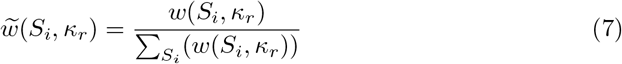

This calculation represents how much each initial condition contributed to the mean 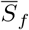 . We found starkly different results for *N*_*max*_ = 9 and 1025 cell simulations (Figs. 4 b and d). For *N*_*max*_ = 9, since the total number of cells is relatively small, initial conditions with *S*_*i*_ *>* 0.5 are common (Fig. 4a). As a result, a wide range of initial conditions contribute substantially to the final abundance of slow killer cells (Fig. 4b). For small environments, we find that relatively rare initial conditions with large *S*_*i*_ can have a substantial contribution. This observation stems from the fact that the slow killer completely is likely to eliminate the fast killer in such conditions, leading to a lottery effect where unlikely initial conditions result in out-sized gains. In particular, for simulations with low relative killing effectiveness *κ*_*r*_, slow killers only survive if the initial population *S*_*i*_ is large, so these large *S*_*i*_ initial conditions contribute substantially, despite their rarity. Conversely, in large spaces, the probability of *S*_*i*_ values significantly larger than 0.5 is very small ((Fig. 4c), so such starting conditions have negligibly small effects.

### 3.2 Effect of system size on invasion

The above analyses show that finite size effects enable slow killing cells to survive at higher rates in small environments than they do in large environments. However, the relative abundance of the slow killer decreases in all conditions studied in Fig. 2, as both strains have the same relative abundance in the planktonic phase and have equal growth rates. We now will focus on the impact of community size when one strain has a numerical advantage over the other strain. These simulations followed a similar protocol as those discussed above, but with different *p*_*i,S*_ values (equation 5, Fig. 5 a). We found that slow killer cells did in fact survive in more conditions, i.e., for a wider range of both *κ*_*r*_ and *p*_*i,S*_, in small environments than in large environments (Fig. 5b and 5c). Surprisingly, we found that stochastic fluctuations even help the slow killer survive in small spaces when *p*_*i,S*_ *<* 0.5, especially if *κ*_*r*_ *>* 0.5. Conversely, slow killers in large spaces are completely eliminated when *p*_*i,S*_ *<* 0.5. For example, if *κ*_*r*_ = 0.37, the slow killer survives in the small environment as long as *p*_*i,S*_ *>* 0.35, but only survives in the large environment if *p*_*i,S*_ *>* 0.6.

**Fig 5.**
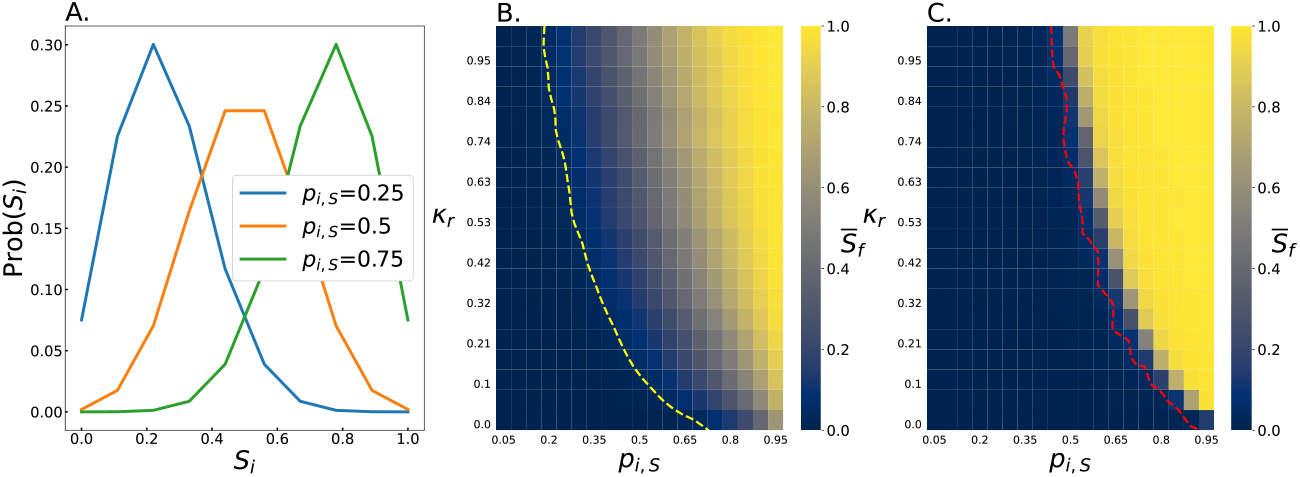
Slow killing cells survive better in small spaces when stochastically seeded from different planktonic abundances. A. The probability distribution of the number of slow cells on a *N*_*max*_ = 9 cell space seeded from planktonic suspensions with relative abundances of slow killers of 0.25, 0.5, and 0.75. The final relative abundance of slow killer cells, 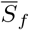, as a function of the relative abundance in the planktonic suspension and the relative killing rate *p*_*i,S*_, *κ*_*r*_. Data are shown for *N*_*max*_ = 9 cell and *N*_*max*_ = 1025 cell simulations (B and C, respectively). The yellow trend line (B) and red trend line (C) show where 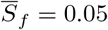. Slow killer cells survive in a much wider range of scenarios in *N*_*max*_ = 9 cell simulations than in *N*_*max*_ = 1025 cell simulations.

We next sought to determine not just the conditions that allow the slow killer to survive, but which conditions actually enable them to increase their relative abundance. We thus plot 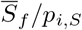 (Fig. 6a and b); when 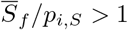, the slow killing strain increased its relative abundance, and when 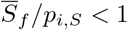, it decreased its relative abundance. In contrast to the above results about survival, we found that the slow killer increases its relative abundance in fewer initial conditions in the small environments than the large environments. However, in large environments 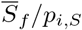 decreases sharply once it drops below 1, such that whatever strain is not favored is almost completely eliminated 6. In contrast, small environments exhibit a smaller range of conditions with 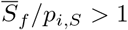 and a larger region of space where 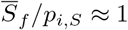. This means that the outcome depends on a combination of the relative killing rates *κ*_*r*_, the invading population size *p*_*i,S*_, and community size.

**Fig 6.**
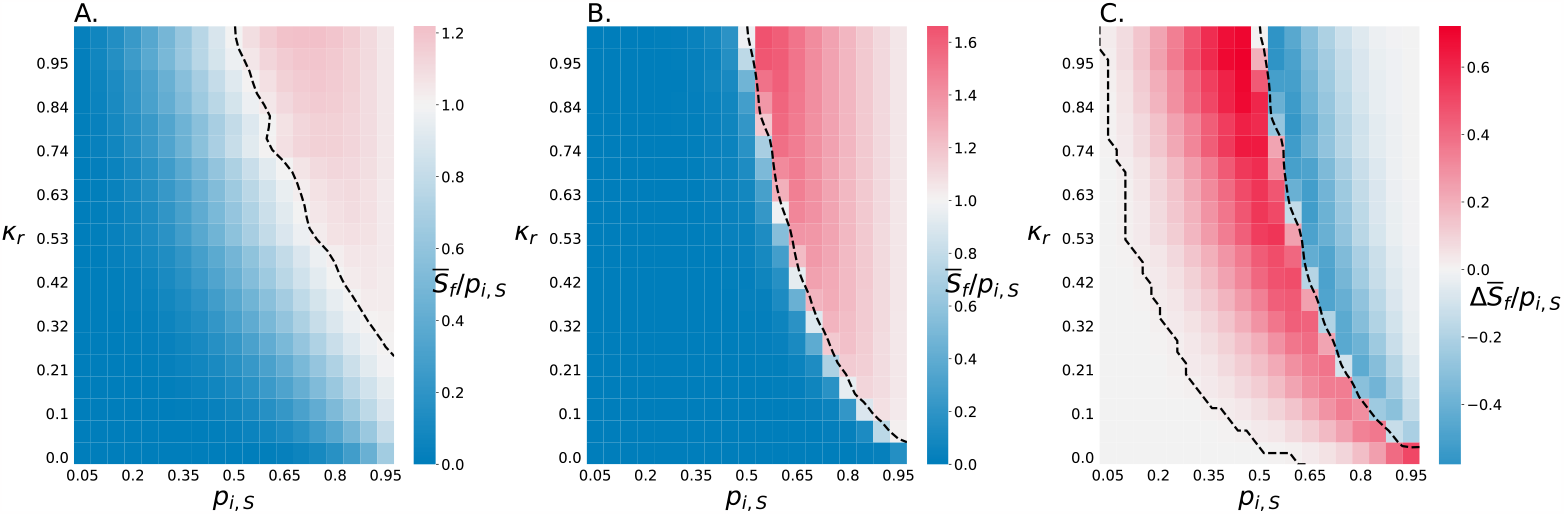
Slow killing cells are more robust against invasion when drastically disadvantaged. A, B. These heat maps plot the final relative abundance of slow killing cells divided by the initial relative abundance of slow killing cells in the planktonic suspension, i.e, 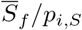, as a function of the relative abundance of slow killers in planktonic suspension and the relative killing rate, *κ*_*r*_, for *N*_*max*_ = 9 cell simulations and *N*_*max*_ = 1025 cell simulations (A and B, respectively). The black trend lines show where 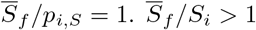 indicates the slow killing strain increases its relative abundance. C. This heat map shows the difference between 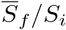 for the 9 and *N*_*max*_ = 1025 cell simulations, i.e., it shows the results from A minus the results from B. Red indicates conditions in which the slow killing strain performs better in the *N*_*max*_ = 9 cell environment, and blue indicates conditions in which the slow killing strain performs better in the *N*_*max*_ = 1025 cell environment. White space indicates there is no difference between the two environments. The black contour line shows where the difference is equal to 0 indicating there is no difference between the large and small space.

**Fig 7.**
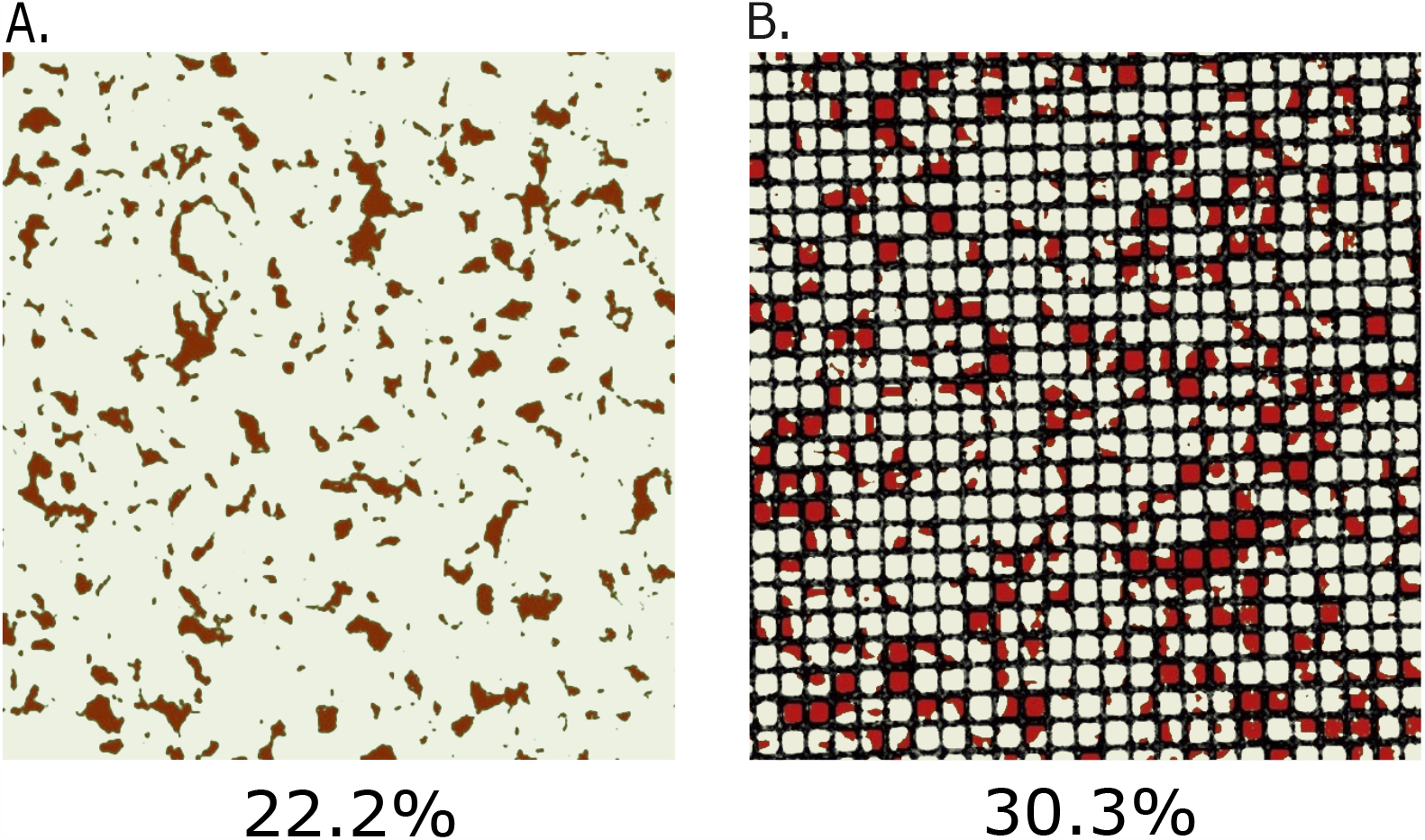
Experiments validate that slow killing strains perform better in small environments. A. Two mutual killing strains of *V. cholerae* were grown for 6 hours in colonies ∼ 1*mm* in diameter on bare agar. The red strain is the slow killer and the white strain is the fast killer. B. The same strains were grown for 6 hours confined to 7.5*μm* square holes in TEM grids. Mean is a result of 5 and 6 different results respectively, p value=0.013.

**Fig 8.**
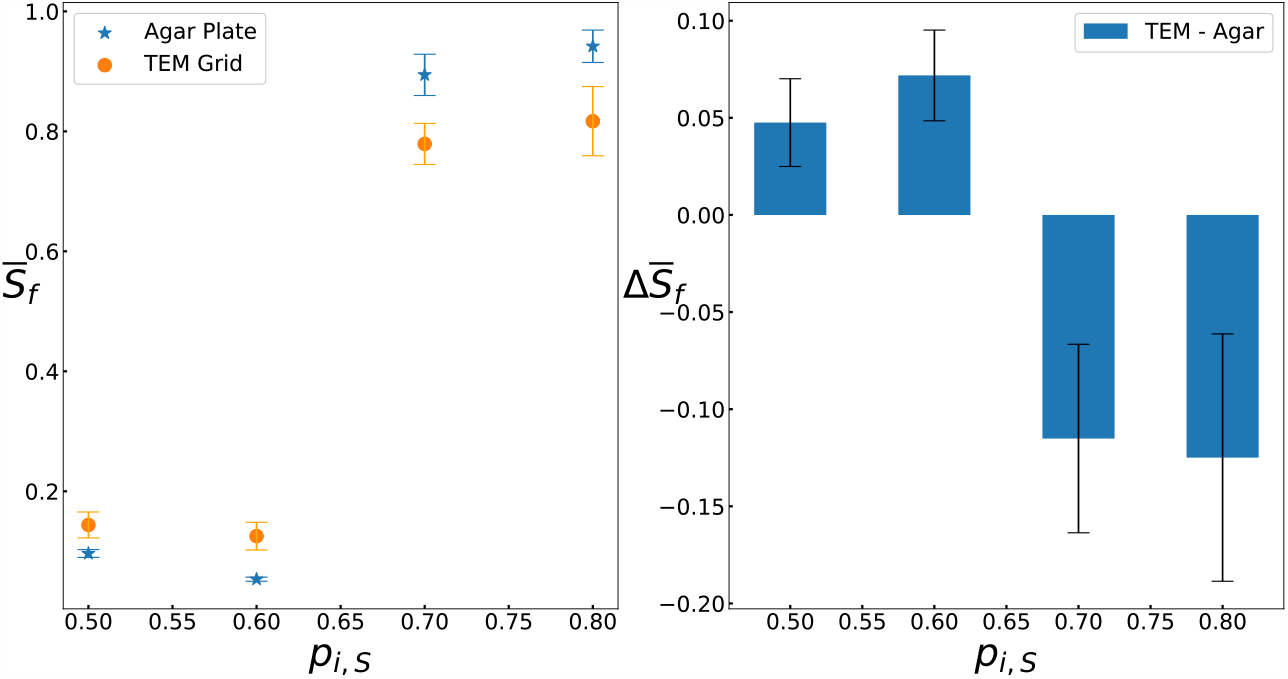
Experiments validate that small spaces decrease fitness difference across different abundances. A. Two mutual killing strains of *V. cholerae* were grown for 8 hours in colonies ∼ 1*mm* in diameter on bare agar and in 7.5*μm* square holes in TEM grids with different starting abundances (*p*_*i,S*_). The means reported are the results of 3,3,6,5 experiments for the open space and 4,3,3,3 for grided.

**Fig 9.**
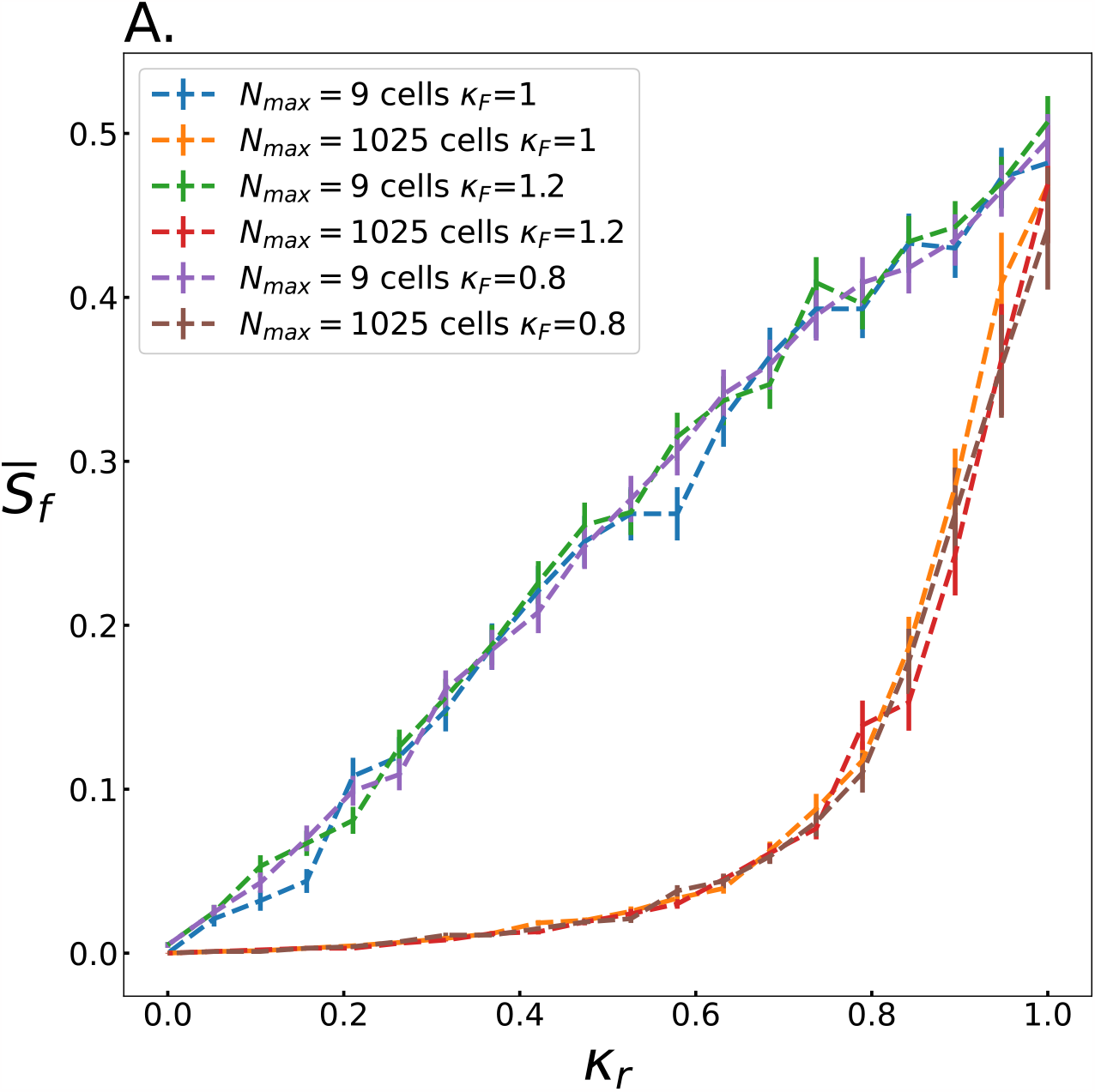
Finite size effects are independent of *κ*_*F*_.

To further explore the impact of community size, we plotted 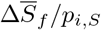, the difference in 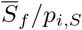 between small and large environments with identical *p*_*i,S*_ (Fig. 6 c). The same contour that separates where the slow killing strain is less fit than the fast killing strain in the large space (see 6 b) separates where the slow killer does better in the small and large environments (Figure 6 c). This quantification makes it clear that the slow killing strain survives better in small spaces with conditions that cause it to lose in the large spaces. This is true outside of some edge cases where the slow killing strain starts with too small *p*_*i,s*_ or *κ*_*r*_ where it does not survive at all. Further, for any condition in which the slow killing strain increases its abundance in the large space, it will increase its own abundance less in the small space.

### 3.3 Experimental validation

The above simulations demonstrate the potential impact of finite size effects on microbial warfare. We now empirically test if these effects can be observed in experiments by growing two strains of mutual killing bacteria in large and small spaces.

Similar to the simulations we describe above, we found that when the competition begins with equal numbers, the slow killing strain ends with larger relative abundance when confined within the TEM grids than when competing on bare agar (0.30 and 0.22, respectively, *p* = 0.013). In fact, the slow killer strain performs 36% better in confinement than out of confinement. These observations support the idea that stochastic fluctuations in small environments decrease the benefit of contact killing speed. The proportion of strains was measured 6 hours after inoculation, sufficient time for T6SS killing to have ceased [9].

We next performed experiments with different initial relative abundances.

Specifically, the slow killer began with relative abundances of *p*_*i,S*_ = 0.5, 0.6, 0.7, and 0.8. We allowed more time to pass (8 hours) before taking measurements across the different starting conditions so as to not compare steady state conditions with dynamic ones given that different starting conditions take different amounts of time to reach a steady state [9]. Similar to our simulations, we found that while the slow killer loses significantly when *p*_*i,S*_ = 0.5 and 0.6, it wins when *p*_*i,S*_ = 0.7 and 0.8, and this is true in large and small spaces. Further, the strain that loses in the large space always does better in the small space. When *p*_*i,S*_ = 0.5 and 0.6, the difference 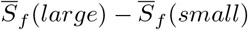 is −0.048 ± 0.023 and −0.072 ± 0.023 (mean ± standard error), respectively, indicating that the slow killer has a larger relative abundance in the small environment. When *p*_*i,S*_ = 0.7 and 0.8, 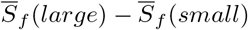 is −0.115 ± 0.049 and −0.125 ± 0.064, respectively, indicating that the slow killer has a smaller relative abundance in the small environment.

## 4 Discussion

Here, we demonstrated that finite size effects can substantially impact the outcome of antagonistic competitions between microbes. In particular, slow contact killing strains survive at a higher rate in smaller spaces than in larger spaces when the two strains initially have equal abundances. We found that in small spaces the final proportion of the slow killing strain is linearly proportional to the ratio of killing effectiveness; this trend persists down to small values of *κ*_*r*_. Conversely, in large spaces the final proportion of slow killing cells decreases rapidly as *κ*_*r*_ decreases from 1.0, and is essentially zero for *κ*_*r*_ *<* 0.4. We found that stochastic fluctuations in the initial seeding of these competitions is the primary source of these effects. Further, we found that slow killing cells survive in more conditions when the initial abundance is varied, but the slow killing strain is more likely to increase its abundance in larger spaces. Finally, we performed experiments to validate these simulations by confining bacteria within TEM grids. We observed the same trend in experiments as we did in simulations; the slow killing strain performed much better in confinement than on bare agar when the initial proportions are equal. Thus, these results demonstrate that finite size effects substantially alter antagonistic competitions between bacteria.

The observation that slow killing strains perform better in small environments than in large environments may also have implications for the evolution of microbial warfare. In particular, studies have found both a significant proportion of bacterial strains have contact killing mechanisms [8], and antagonistic interactions are commonplace [1], but there also exists a great diversity of killing ability [11]. These observations seem to indicate that bacteria do not experience an unmitigated arms race in which only the best killing strains can survive. The results presented here suggest that competition in small spaces decreases the value of contact killing speed, which may contribute to the subversion of this arms race. Crucially, when the slow killer has a numerical advantage and “wins” in the large environment, it does not perform as well in the small environment. Finite size effects may mitigate differences between strains in general. Such effects would decrease the fitness advantage of the “superior” strain, constraining its ability to takeover its environment. As a result, finite size effects may lead to more diversity in killing ability.

There are many known mechanisms that limit the utility of microbial antagonism. For example, many studies have found that extracellular matrix production can prevent killing [37]. Rock-paper-scissors game theory can lead to antagonism being a poor strategy. Further, strains can evolve various resistance mechanisms [38–41]. In a different vein, a toxin may not be effective against a novel competitor. However, all of these mechanisms have to do with the behavior of bacteria and their competitors. Here, we highlight that spatial constraints size—a factor that bacteria cannot control—can similarly constrain the benefits of microbial antagonism.

Previous publications found that dead cell debris accumulates at the interface between contact killing strains [9, 10], eventually preventing contact killing from occurring. This effect may limit the impact of finite size effects in nature. In large environments, slow killers will do better as eventually dead cell debris will prevent further killing, enabling their survival. In small environments, dead cell debris may not accumulate quickly enough to limit contact killing, but when it does it may limit access to “jackpots,” i.e., a single strain completely taking over its environment. However, our experiments and simulations exhibit roughly similar effect sizes (though comparisons are imperfect), suggesting that the finite size effects impact outcomes before dead cell debris prevents contact killing. Further study is necessary to determine which strain, the fast or slow killer, would receive the most benefit in this scenario. Nonetheless, as demonstrated experimentally here, finite size effects do still impact the outcomes of experiments in confinement.

Details about how cells attach to surfaces may play a large role in determining the impact of finite size effects. We model attachment as occurring randomly where each cell has some chance of being either strain and that chance is independent of the cells around it as we assume a well-mixed suspension. However, microbes in nature may form aggregates, which then attach to the surface with a pre-existing spatial structure [42, 43]. These aggregates can be as large as 100*um* which could enhance the impact of finite size effects, as aggregates would increase the size of fluctuations in the initial abundance of a given strain [44]. Conversely, clumping with close relative cells will be less likely to impact the distribution of cells in large systems. In a related vein, while we explored scenarios where many cells attach simultaneously, in nature cells can attach to a surface at different times. This scenario would likely also accentuate finite size effects, as slow killer cells that attach first may have the chance to reproduce before competitors attach. In other words, there may be priority effects that further enhance finite size effects.

Microbes often live in small environments in nature. While the exact proportion of large and small system sizes in nature would be exceedingly difficult to quantify, large open spaces that do not experience significant shear forces are likely exceptional. Thus, understanding how microbial warfare proceeds in nature necessitates understanding how it proceeds in small environments. The results here suggest that differences in fitness between slow and fast killing strains, as well as rare and common strains, are reduced in small environments. Thus, small environments may lead to greater diversity both because of their smaller effective population size as well as because fitness differences are reduced. Future work may try to generalize these results beyond microbial antagonism.

## 5 Supporting information

It is possible that the finite size effects observed were dependent on the fast killing strain’s killing rate *κ*_*F*_ = 1. In order to discern if this was the case, we ran simulations were *κ*_*F*_ = 0.8 and *κ*_*F*_ = 1.2 in the *L* = 3 and *L* = 31 spaces and recorded 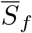 similar to 2.

We found that changing *κ*_*F*_ had no effect in either the small or large environments9. The trend lines for all values of *κ*_*F*_ followed the same trajectory where the small environments produced a linear relationship between 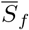 and *κ*_*r*_ from *κ*_*r*_ = 0 to *κ*_*r*_ = 1 while the large environments led to no survival of the slow killing strain below *κ*_*r*_ *<* 0.4.

## References

1. Rendueles O, Ghigo JM. Mechanisms of Competition in Biofilm Communities. In: Microbial Biofilms. John Wiley &Sons, Ltd; 2015. p. 319–342. Available from: https://onlinelibrary.wiley.com/doi/abs/10.1128/9781555817466.ch16.

2. Madsen JS, Røder HL, Russel J, Sørensen H, Burmølle M, Sørensen SJ. Coexistence facilitates interspecific biofilm formation in complex microbial communities. Environmental Microbiology. 2016;18(8):2565–2574. doi:10.1111/1462-2920.13335.

3. Flemming HC, Wuertz S. Bacteria and archaea on Earth and their abundance in biofilms. Nature Reviews Microbiology. 2019;17(4):247–260. doi:10.1038/s41579-019-0158-9.

4. Li YH, Tian X. Quorum Sensing and Bacterial Social Interactions in Biofilms. Sensors. 2012;12(3):2519–2538. doi:10.3390/s120302519.

5. Blanchard AE, Lu T. Bacterial social interactions drive the emergence of differential spatial colony structures. BMC Systems Biology. 2015;9(1):59. doi:10.1186/s12918-015-0188-5.

6. Yang L, Liu Y, Wu H, Høiby N, Molin S, Song Zj. Current understanding of multispecies biofilms. International Journal of Oral Science. 2011;3(2):74–81. doi:10.4248/IJOS11027.

7. Granato ET, Meiller-Legrand TA, Foster KR. The Evolution and Ecology of Bacterial Warfare. Current Biology. 2019;29(11):R521–R537. doi:10.1016/j.cub.2019.04.024.

8. Ho BT, Dong TG, Mekalanos JJ. A View to a Kill: The Bacterial Type VI Secretion System. Cell Host &Microbe. 2014;15(1):9–21. doi:10.1016/j.chom.2013.11.008.

9. Steinbach G, Crisan C, Ng SL, Hammer BK, Yunker PJ. Accumulation of dead cells from contact killing facilitates coexistence in bacterial biofilms. Journal of The Royal Society Interface. 2020;17(173):20200486. doi:10.1098/rsif.2020.0486.

10. Smith WPJ, Vettiger A, Winter J, Ryser T, Comstock LE, Basler M, et al. The evolution of the type VI secretion system as a disintegration weapon. PLOS Biology. 2020;18(5):e3000720. doi:10.1371/journal.pbio.3000720.

11. Bernardy EE, Turnsek MA, Wilson SK, Tarr CL, Hammer BK. Diversity of Clinical and Environmental Isolates of Vibrio cholerae in Natural Transformation and Contact-Dependent Bacterial Killing Indicative of Type VI Secretion System Activity. Applied and Environmental Microbiology. 2016;82(9):2833–2842. doi:10.1128/AEM.00351-16.

12. Breen P, Winters AD, Theis KR, Withey JH. The Vibrio cholerae Type Six Secretion System Is Dispensable for Colonization but Affects Pathogenesis and the Structure of Zebrafish Intestinal Microbiome. Infection and Immunity. 2021;89(9):e00151–21. doi:10.1128/IAI.00151-21.

13. Lucero CT, Lorda GS, Luduenã LM, Nievas F, Bogino PC, Angelini J, et al. Participation of type VI secretion system in plant colonization of phosphate solubilizing bacteria. Rhizosphere. 2022;24:100582. doi:10.1016/j.rhisph.2022.100582.

14. Nadell CD, Drescher K, Foster KR. Spatial structure, cooperation and competition in biofilms. Nature Reviews Microbiology. 2016;14(9):589–600. doi:10.1038/nrmicro.2016.84.

15. McNally L, Bernardy E, Thomas J, Kalziqi A, Pentz J, Brown SP, et al. Killing by Type VI secretion drives genetic phase separation and correlates with increased cooperation. Nature Communications. 2017;8(1):14371. doi:10.1038/ncomms14371.

16. Yanni D, Márquez-Zacarías P, Yunker PJ, Ratcliff WC. Drivers of Spatial Structure in Social Microbial Communities. Current Biology. 2019;29(11):R545–R550. doi:10.1016/j.cub.2019.03.068.

17. Hol FJH, Rotem O, Jurkevitch E, Dekker C, Koster DA. Bacterial predator–prey dynamics in microscale patchy landscapes. Proceedings of the Royal Society B: Biological Sciences. 2016;283(1824):20152154. doi:10.1098/rspb.2015.2154.

18. Deschaine BM, Heysel AR, Lenhart BA, Murphy HA. Biofilm formation and toxin production provide a fitness advantage in mixed colonies of environmental yeast isolates. Ecology and Evolution. 2018;8(11):5541–5550. doi:10.1002/ece3.4082.

19. Ursell T. Structured environments foster competitor coexistence by manipulating interspecies interfaces. PLOS Computational Biology. 2021;17(1):e1007762. doi:10.1371/journal.pcbi.1007762.

20. Vallespir Lowery N, Ursell T. Structured environments fundamentally alter dynamics and stability of ecological communities. Proceedings of the National Academy of Sciences. 2019;116(2):379–388. doi:10.1073/pnas.1811887116.

21. Brandwein M, Steinberg D, Meshner S. Microbial biofilms and the human skin microbiome. npj Biofilms and Microbiomes. 2016;2(1):1–6. doi:10.1038/s41522-016-0004-z.

22. Wielen PWJJ, Keuzenkamp DA, Lipman LJA, Knapen F, Biesterveld S. Spatial and Temporal Variation of the Intestinal Bacterial Community in Commercially Raised Broiler Chickens During Growth. Microbial Ecology. 2002;44(3):286–293. doi:10.1007/s00248-002-2015-y.

23. Probandt D, Eickhorst T, Ellrott A, Amann R, Knittel K. Microbial life on a sand grain: from bulk sediment to single grains. The ISME Journal. 2018;12(2):623–633. doi:10.1038/ismej.2017.197.

24. Kim MK, Drescher K, Pak OS, Bassler BL, Stone HA. Filaments in curved streamlines: rapid formation of Staphylococcus aureus biofilm streamers. New Journal of Physics. 2014;16(6):065024. doi:10.1088/1367-2630/16/6/065024.

25. Connell JL, Wessel AK, Parsek MR, Ellington AD, Whiteley M, Shear JB. Probing Prokaryotic Social Behaviors with Bacterial “Lobster Traps”. mBio. 2010;1(4):e00202–10. doi:10.1128/mBio.00202-10.

26. Hartmann R, Singh PK, Pearce P, Mok R, Song B, Díaz-Pascual F, et al. Emergence of three-dimensional order and structure in growing biofilms. Nature Physics. 2019;15(3):251–256. doi:10.1038/s41567-018-0356-9.

27. Gniewek P, Schreck CF, Hallatschek O. Biomechanical Feedback Strengthens Jammed Cellular Packings. Physical Review Letters. 2019;122(20):208102. doi:10.1103/PhysRevLett.122.208102.

28. Jacobeen S, Pentz JT, Graba EC, Brandys CG, Ratcliff WC, Yunker PJ. Cellular packing, mechanical stress and the evolution of multicellularity. Nature Physics. 2018;14(3):286–290. doi:10.1038/s41567-017-0002-y.

29. Delarue M, Hartung J, Schreck C, Gniewek P, Hu L, Herminghaus S, et al. Self-driven jamming in growing microbial populations. Nature Physics. 2016;12(8):762–766. doi:10.1038/nphys3741.

30. Schreck CF, O‘Hern CS, Silbert LE. Tuning jammed frictionless disk packings from isostatic to hyperstatic. Physical Review E. 2011;84(1):011305. doi:10.1103/PhysRevE.84.011305.

31. O ‘Hern CS, Silbert LE, Liu AJ, Nagel SR. Jamming at zero temperature and zero applied stress: The epitome of disorder. Physical Review E. 2003;68(1):011306. doi:10.1103/PhysRevE.68.011306.

32. Cheah SE, Li J, Nation RL, Bulitta JB. Novel Rate-Area-Shape Modeling Approach To Quantify Bacterial Killing and Regrowth for In Vitro Static Time-Kill Studies. Antimicrobial Agents and Chemotherapy. 2014;59(1):381–388. doi:10.1128/AAC.04182-14.

33. Thomas J, Watve SS, Ratcliff WC, Hammer BK. Horizontal Gene Transfer of Functional Type VI Killing Genes by Natural Transformation. mBio. 2017;8(4):e00654–17. doi:10.1128/mBio.00654-17.

34. Boyer F, Fichant G, Berthod J, Vandenbrouck Y, Attree I. Dissecting the bacterial type VI secretion system by a genome wide in silico analysis: what can be learned from available microbial genomic resources? BMC Genomics. 2009;10(1):104. doi:10.1186/1471-2164-10-104.

35. Venkateswaran K. VIBRIO Standard Cultural Methods and Molecular Detection Techniques in Foods. In: Robinson RK, editor. Encyclopedia of Food Microbiology. Oxford: Elsevier; 1999. p. 2248–2258. Available from: https://www.sciencedirect.com/science/article/pii/B0122270703016652.

36. Otsu N. A Threshold Selection Method from Gray-Level Histograms. IEEE Transactions on Systems, Man, and Cybernetics. 1979;9(1):62–66. doi:10.1109/TSMC.1979.4310076.

37. Teschler JK, Jiménez-Siebert E, Jeckel H, Singh PK, Park JH, Pukatzki S, et al. VxrB Influences Antagonism within Biofilms by Controlling Competition through Extracellular Matrix Production and Type 6 Secretion. mBio. 2022;13(4):e01885–22. doi:10.1128/mbio.01885-22.

38. MacGillivray KA, Ng SL, Wiesenfeld S, Guest RL, Jubery T, Silhavy TJ, et al. Trade-offs constrain adaptive pathways to T6 survival. bioRxiv. 2022;doi:10.1101/2022.09.02.506412.

39. Crisan CV, Nichols HL, Wiesenfeld S, Steinbach G, Yunker PJ, Hammer BK. Glucose confers protection to Escherichia coli against contact killing by Vibrio cholerae. Scientific Reports. 2021;11(1):2935. doi:10.1038/s41598-021-81813-4.

40. Toska J, Ho BT, Mekalanos JJ. Exopolysaccharide protects Vibrio cholerae from exogenous attacks by the type 6 secretion system. Proceedings of the National Academy of Sciences of the United States of America. 2018;115(31):7997–8002. doi:10.1073/pnas.1808469115.

41. Granato ET, Smith WPJ, Foster KR. Collective protection against the type VI secretion system in bacteria. bioRxiv. 2022;doi:10.1101/2022.09.12.507624.

42. Trunk T, Khalil HS, Leo JC. Bacterial autoaggregation. AIMS Microbiology. 2018;4(1):140–164. doi:10.3934/microbiol.2018.1.140.

43. Sauer K, Stoodley P, Goeres DM, Hall-Stoodley L, Burmølle M, Stewart PS, et al. The biofilm life cycle: expanding the conceptual model of biofilm formation. Nature Reviews Microbiology. 2022;20(10):608–620. doi:10.1038/s41579-022-00767-0.

44. Geng J, Henry N. Short time-scale bacterial adhesion dynamics. Bacterial Adhesion: Chemistry, Biology and Physics. 2011; p. 315–331.

